# Penetrance of Parkinson’s disease in *LRRK2* p.G2019S carriers is modified by a polygenic risk score

**DOI:** 10.1101/738260

**Authors:** Hirotaka Iwaki, Cornelis Blauwendraat, Mary B. Makarious, Sara Bandrés-Ciga, Hampton L. Leonard, J. Raphael Gibbs, Dena G. Hernandez, Sonia W. Scholz, Faraz Faghri, International Parkinson’s Disease Genomics Consortium (IPDGC), Mike A. Nalls, Andrew B. Singleton

## Abstract

**Background:** While the LRRK2 p.G2019S mutation has been demonstrated to be a strong risk factor for Parkinson’s Disease (PD), factors that contribute to penetrance among carriers, other than aging, have not been well identified.

**Objectives:** To evaluate whether a cumulative genetic risk identified in the recent genome-wide study is associated with penetrance of PD among p.G2019S mutation carriers.

**Methods:** We included p.G2019S heterozygote carriers with European ancestry in three genetic cohorts in which the mutation carriers with and without PD were selectively recruited. We also included the carriers from two datasets: one from a case-control setting without selection of mutation carriers, and the other from a population sampling. The associations between PRS constructed from 89 variants reported in Nalls et al. and PD were tested and meta-analyzed. We also explored the interaction of age and PRS.

**Results:** After excluding 8 homozygotes, 833 p.G2019S heterozygote carriers (439 PD and 394 unaffected) were analyzed. PRS was associated with a higher penetrance of PD (OR 1.34, 95% C.I. [1.09, 1.64] per +1 SD, *P* = 0.005). In addition, associations with PRS and penetrance were stronger in the younger participants (main effect: OR 1.28 [1.04, 1.58] per +1 SD, *P* = 0.022; interaction effect: OR 0.78 [0.64, 0.94] per +1 SD and +10 years of age, *P* = 0.008).

**Conclusions:** Our results suggest that there is a genetic contribution for penetrance of PD among p.G2019S carriers. These results have important etiologic consequences and potential impact on the selection of subjects for clinical trials.

## Introduction

Parkinson’s disease (PD) is a complex genetic disorder, where rare and highly damaging variants as well as common risk variants play a role in its etiology.^1^ The *LRRK2* p.G2019S (rs34637584) mutation is one of the major known contributors to PD.^2,3^ The p.G2019S mutation has an estimated prevalence of approximately 1% in the PD population of European ancestry, with much higher frequencies being reported for North African Berber Arab populations and European subpopulations with high Ashkenazi Jewish ancestry.^4–7^ The p.G2019S mutation is not fully penetrant, with risk of PD for carriers increasing with age. At age 80 years, 25% to 42.5% of carriers will have PD.^7,8^ Critical question that remain centers on what the determinants of p.G2019S penetrance are. Previously, it has been reported that a variant in *DNM3*, encoding the vesicular transport protein dynamin 3, is a potential *LRRK2* p.G2019S age-at-onset modifier, lowering the age at onset by approximately 8 years.^9^ However, other reports have not replicated this finding and instead nominated variants in *SNCA* as modifiers of *LRRK2* mutation penetrance.^10^ The underlying pathogenic mechanism of *LRRK2*-linked PD is currently unknown, but a great deal of focus has been placed on the ability of the p.G2019S mutation to increase LRRK2 kinase activity, especially since loss-of-function mutations do not seem to be contributing to disease.^11,12^ In the last several years, genome-wide association studies (GWAS) have identified an ever-increasing number of risk loci for PD. Individually, each of these loci confers modest effects on risk for disease. Polygenic risk score (PRS) represents known cumulative genetic risk across these loci in each assayed individual. PRS reveals that, collectively, these risk variants confer considerable risk for disease, with those in the top decile of genetic risk being sixfold more likely to have PD than those in the lowest decile of genetic risk in the European population.^1^ PRS is also highly correlated with age at onset of PD,^13,14^ and is associated with penetrance of damaging *GBA* variants.^15^ Here, we investigate the influence of the latest PD PRS on age at onset and penetrance of PD in p.G2019S carriers using several large cohorts.

## Methods

### Whole genome sequencing

Genetics and clinical data used in the preparation of this article were obtained from the Parkinson’s Progression Markers Initiative (PPMI) database (www.ppmi-info.org/data). (For up-to-date information on the PPMI study, visit www.ppmi-info.org.) In this study, we included three genetic cohorts: Parkinson’s Progression Markers Initiative Genetic Cohort (PPMI_GC), Parkinson’s Progression Markers Initiative Genetic Registry (PPMI_GR), and the LRRK2 Consortium Cohort (LCC), hereafter collectively named the *Genetic Cohort Dataset*. These cohorts selectively included people with specific high-risk genetic variants, such as carrying p.G2019S, for both cases and unaffected individuals. Whole genome sequencing (WGS) data was generated at the Laboratory of Neurogenetics (LNG) at the National Institutes of Health, and detailed methods are available from the study website. After downloading, subsequent genotypes were filtered to exclude minor allele frequency of less than 0.01, missing rate more than 5%, and the test for Hardy–Weinberg equilibrium threshold of 1.0E-4 in controls. Following quality control, 185 carriers of p.G2019S (PD/Non-PD = 88/97) in PPMI_GC, 75 (38/37) in PPMI_GR, and 175 (107/68) in LCC were included in the analysis. Among them, 7 individuals (3 in PPMI_GC, 3 in PPMI_GR, and 1 in LCC, all cases) were homozygotes.

### Genotype data from the International Parkinson Disease Genomics Consortium

Aggregated genotype data was obtained from the International Parkinson Disease Genomics Consortium (IPDGC), as previously described.^1^ In brief, genotypes were processed using standard pipelines and imputed using the HRC imputation panel r1.1 2016^16^ via the Michigan imputation server under the default setting with phasing using the EAGLE option.^17^ Genotypes were filtered for imputation quality *R*^2^ > 0.8. Following quality control criteria similar to that applied to the Genetic Cohort Dataset, a total of 227 carriers were included in the analysis, of which 208 were PD cases and 19 were controls. Among them, one PD patient was homozygous. For composing PRS, we used a threshold of *R*^2^ > 0.3 to obtain dosages of more risk-associated alleles because only 64 variants were passed with the cutoff of 0.8 in *R*^2^.

### UK Biobank data

The United Kingdom (UK) Biobank (UKBB) is a large, long-term biobank study in the UK that includes genetic data and a wide range of phenotypes of approximately 500,000 individuals.^18^ PD case-control status was based on multiple field codes, including 41202, 41204, 40002, and 20002. *LRRK2* p.G2019S was directly genotyped in the UK Biobank cohort (rs34637584, *n* = 314), and the concordance rate in the whole-exome sequencing data^19^ was 100%. After applying the same quality control steps as above, 179 carriers were identified, of which 6 were reported to be PD cases; all were heterozygotes.

### Polygenic risk score calculation

PRS was calculated incorporating the risk loci previously associated with PD.^1^ In the calculation of PRS, risk allele dosages were summed with weights as their published beta estimates, giving greater weight to alleles with higher estimates. Then, PRS were standardized to each cohort level. For the UKBB and IPDGC imputed genotype data, not all 89 variants were available due to lower imputation scores (*R*^2^ > 0.8 for UKBB and *R*^2^ > 0.3 for IPDGC); therefore, 88 variants in UKBB and a median of 89 variants (range: 82–89) in IPDGC were used.

### Statistical analysis

Our primary analysis was to test the association between PRS and PD odds using a fixed effect meta-analysis model among the mutation carriers. First, a logistic regression model was applied for the main effect of PRS for the Genetic Cohort Dataset (PPMI_GC/PPMI_GR/LRRK2), IPDGC dataset, and UKBB dataset separately. The data were adjusted for age, sex, cohort, and PCs representing population structure. Across all datasets, the linear age upon recruitment was used for carriers without PD and the age at diagnosis for carriers with PD, whereas the age at onset was used for the IPDGC dataset and the age at recruitment for the UKBB dataset. The age data were centered—the original value subtracted by the mean age at each cohort—to accommodate square age variable in the later step. To adjust for population structure, PC1–10 were used for the Genetic Cohort Dataset and IPDGC dataset, and PC1–5 were used for the UKBB dataset because of its size.

In the secondary analysis, the interaction between age and PRS for PD odds was tested. In this analysis, we excluded the UKBB dataset because of the small number of PD cases. We meta-analyzed the dataset level ORs for the main effect and the interaction effect between age and PRS as in the similar model above, but we further adjusted for square age in the Genetic Cohort Dataset and IPDGC dataset. Square age was significantly associated with the PD odds in these two datasets; as the UKBB dataset cannot accommodate the variable because of too few PD cases, we thus could not include square age in the primary analysis.

Also, the association of PRS on age at diagnosis (Genetic Cohort Dataset) or age at onset (IPDGC dataset) for the carriers with PD was analyzed in an exploratory manner. Adjusted covariates included age, square age, sex, cohorts, and PC1–10.

There were 8 homozygotes of p.G2019S mutation in the analysis set, and we excluded them from all analyses. Analyses were conducted by using R version 3.5.1 (https://cran.r-project.org/), with a significance level of 0.05 (two-sided). With the sample size here, we had 90% power for the primary analysis and 35% power for the exploratory analysis on age at onset/diagnosis, assuming that the associations were the same as in the previous report conducted in a general case-control setting in the European population.^1^

### Patients Consent

Participants’ information and genetic samples were obtained under appropriate written consent and with local institutional and ethical approvals for all cohorts and datasets in this study.

## Results

### Data overview

Table 1 is a summary of participants with and without the *LRRK2* p.G2019S mutation in the datasets under study. The IPDGC dataset consists of 14,225 cases and 16,543 controls with known *LRRK2* p.G2019S status; only 1.4% of the cases had the *LRRK2* p.G2019S mutation, and 0.11% of the controls carried the mutation. The prevalence of PD in carriers is consistent with results from a previous report of 1% of PD patients carrying the mutation.^7^ Among the 362,884 participants passing filtering criteria in the UKBB, 0.049% were carriers of the *LRRK2* p.G2019S mutation. While the overall prevalence of PD in this dataset was 0.33%, 3.36% of carriers had PD, indicating an approximately 10 times higher crude risk of PD associated with the mutation. This is consistent with the crude relative risk of 13 in UKBB. In total, of 3 datasets there were 8 homozygotes reported as PD cases. They had a higher PRS and older age at diagnosis compared to heterozygous PD cases (Supplemental Table 1), although the differences were not statistically significant. Because of the small number of homozygotes, we excluded them from downstream analyses.

**Table1:**
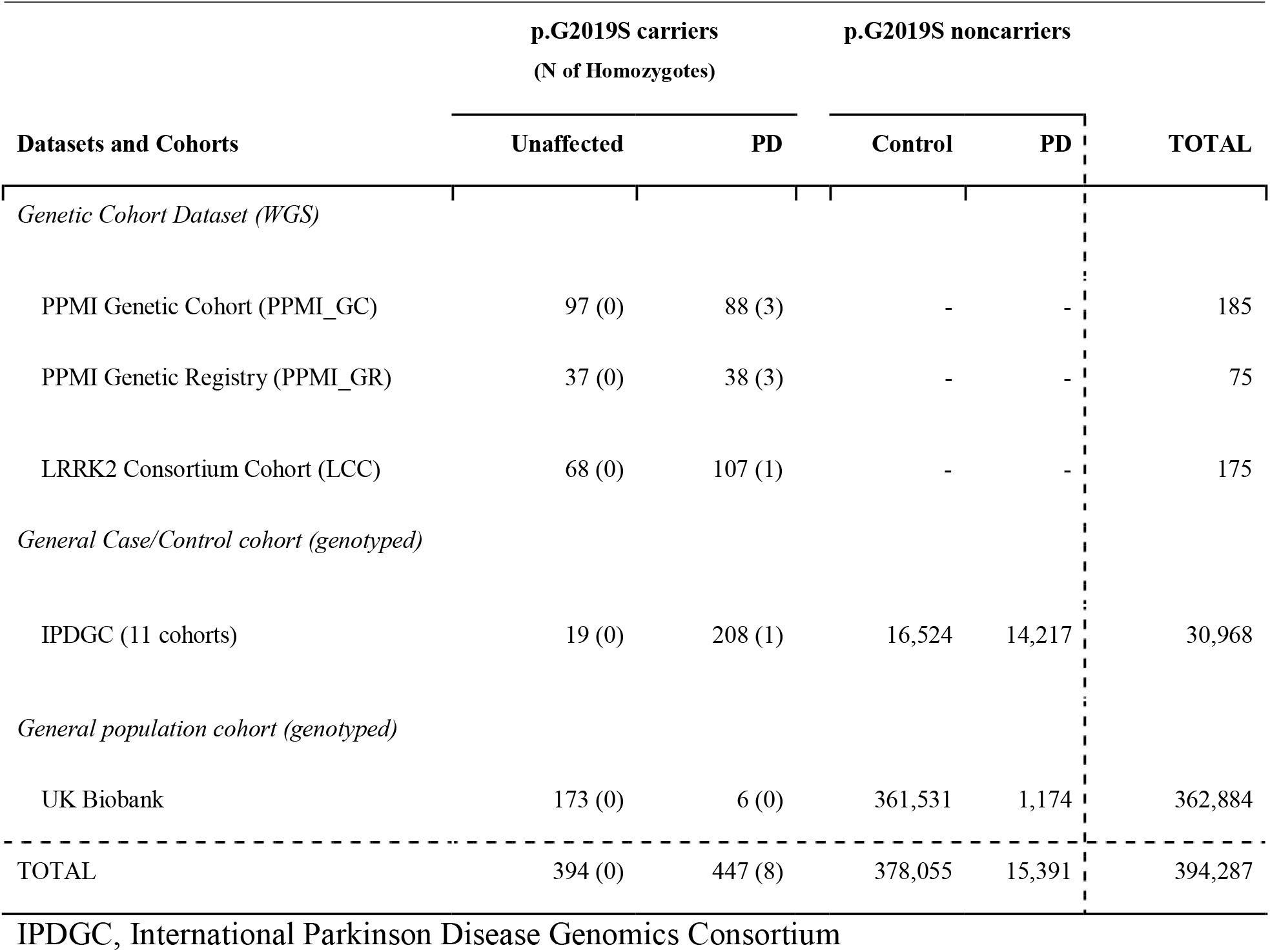
Overview of data.

### Genetic risk score modifies penetrance of LRRK2 p.G2019S

The unadjusted plots of PRS among cases and unaffected individuals in the Genetic Cohort Dataset showed that the mean of the PRS was higher in cases among carriers (Figure 1). The means of PRS in cases were also higher than those in controls among noncarriers in the IPDGC and UKBB datasets (Supplemental Figure 1). Meta-analysis results of the three datasets showed that a higher PRS was significantly associated with a higher odds ratio of PD (OR 1.34, 95% C.I. [1.09, 1.64] per + 1SD [per standard deviation of increase from the cohort mean, *P* = 0.005]; Figure 2A).

**Figure 1:**
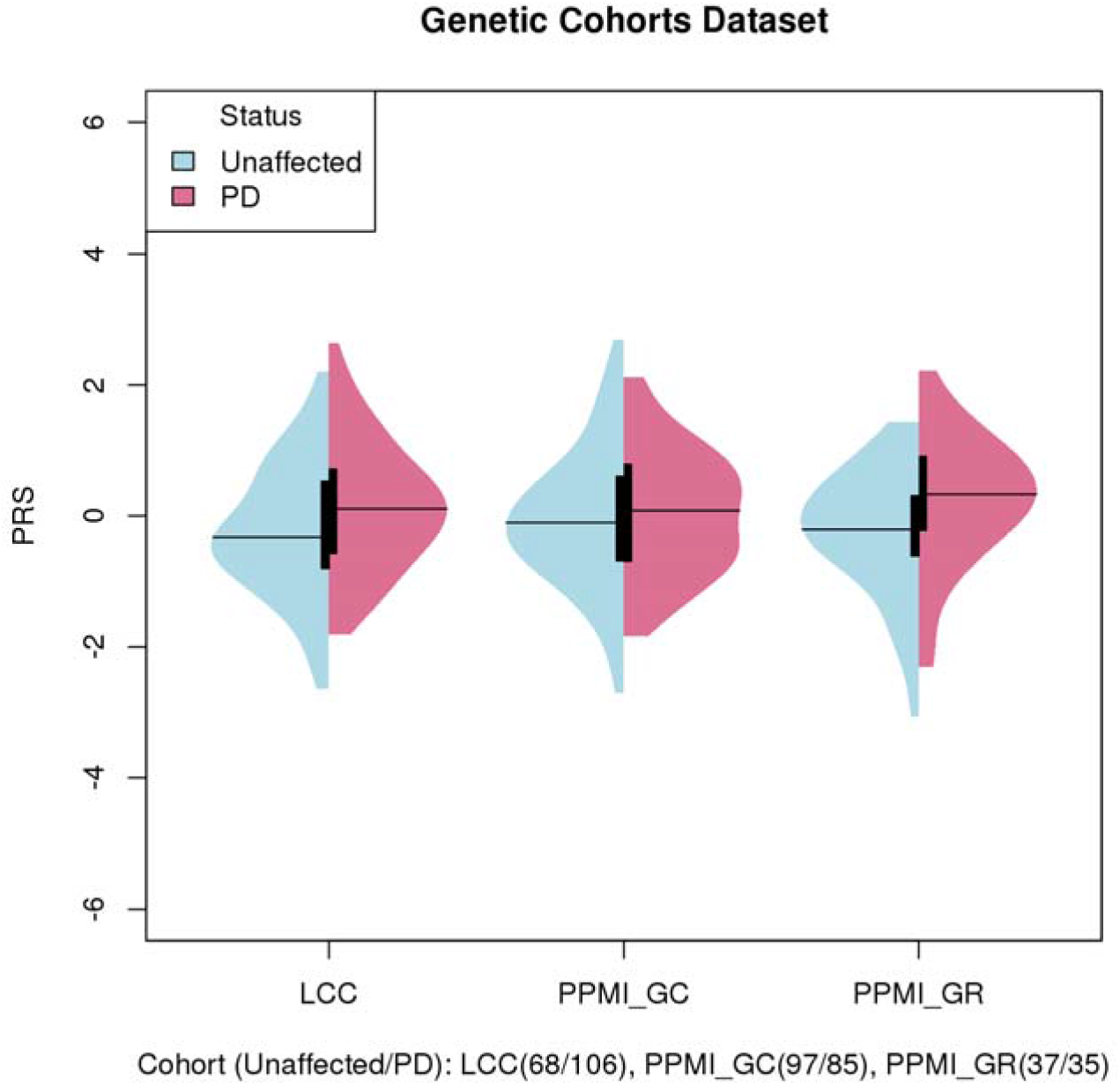
PRS of cases and controls in genetic cohorts. Legend: The means of PRS in cases were higher than unaffected among LRRK2 p.G2019S mutation carriers. LCC, LRRK2 Consortium Cohort; PPMI_GC, Parkinson’s Progression Markers Initiative Genetic Cohort; PPMI_GR, Parkinson’s Progression Markers Initiative Genetic Registry.

**Figure 2:**
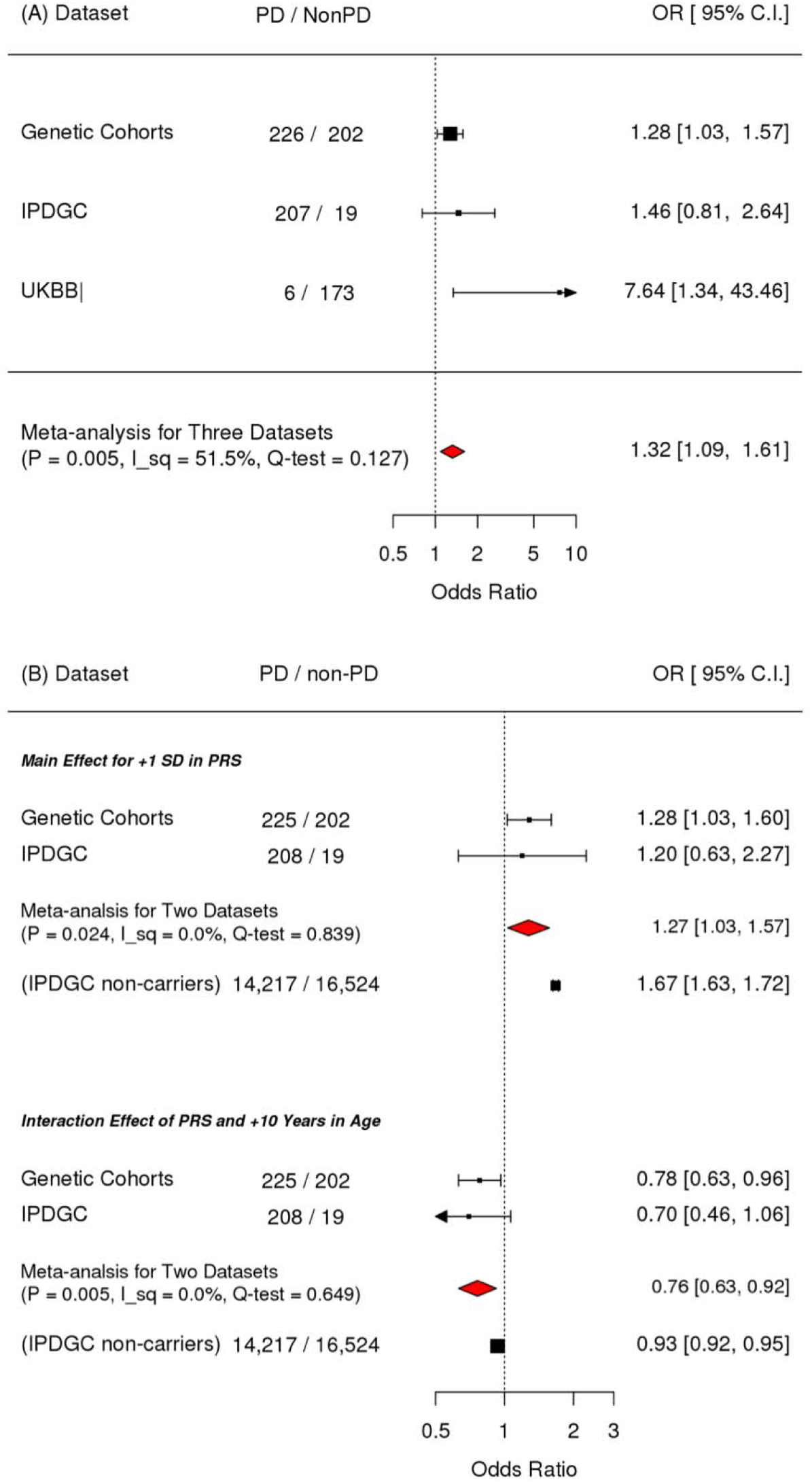
Meta-analysis for PRS, and its interaction with age on penetrance. Legend: A. Primary analysis of testing the association between PRS and PD showed that PRS was significantly associated with the PD odds. B. Secondary analysis model with PRS and the interaction between PRS and age showed that the main effect of PRS as well as the interaction between PRS and age were significantly associated with the odds of PD, but in the opposite direction. Compared to the interaction effect among non-carriers in IPDGC, the magnitude of the interaction among the mutation carriers was large. OR, Odds ratio; I_sq, I square (%); Q-test, P-value for the test of Heterogeneity; IPDGC, International Parkinson Disease Genomics Consortium. Odds ratio were adjusted for study, age, square age, sex and PC1-10

### Stronger association of genetic risk score for PD penetrance in younger carriers

In the secondary analysis examining the interaction between PRS and age, the main effect of PRS as well as the interaction between PRS and age were significantly associated with the odds of PD, but in the opposite direction (main effect: OR 1.28 [1.04, 1.58] per +1 SD, *P* = 0.022; interaction effect: OR 0.78 [0.64, 0.94] per +1 SD and +10 years of age, *P* = 0.008; Figure 2B). This suggests that in the studied cohorts, the magnitude of association between PRS and PD odds were stronger when the age of the participants was younger. We added post-hoc analyses stratified by age (<55 years old, 55-65 years old and 65< years old) to illustrate the stronger association of PRS in younger age in carriers. (Figure 3). The analyses examining the main effect and interaction effect among nonmutation carriers in the IPDGC dataset showed that these effects were also significant in nonmutation carriers (the main term for +1 SD in PRS: OR 1.6, *P* = 2.5E-245; the interaction term of +1 SD in PRS and +10 years in age: 0.94, *P* = 1.4E-9). However, the magnitude of the interaction effect was smaller than that in *LRRK2* p.G2019S carriers, and the magnitudes of associations were significantly different between carriers and non-carriers in the test of homogeneity (P = 0.044).

**Figure 3:**
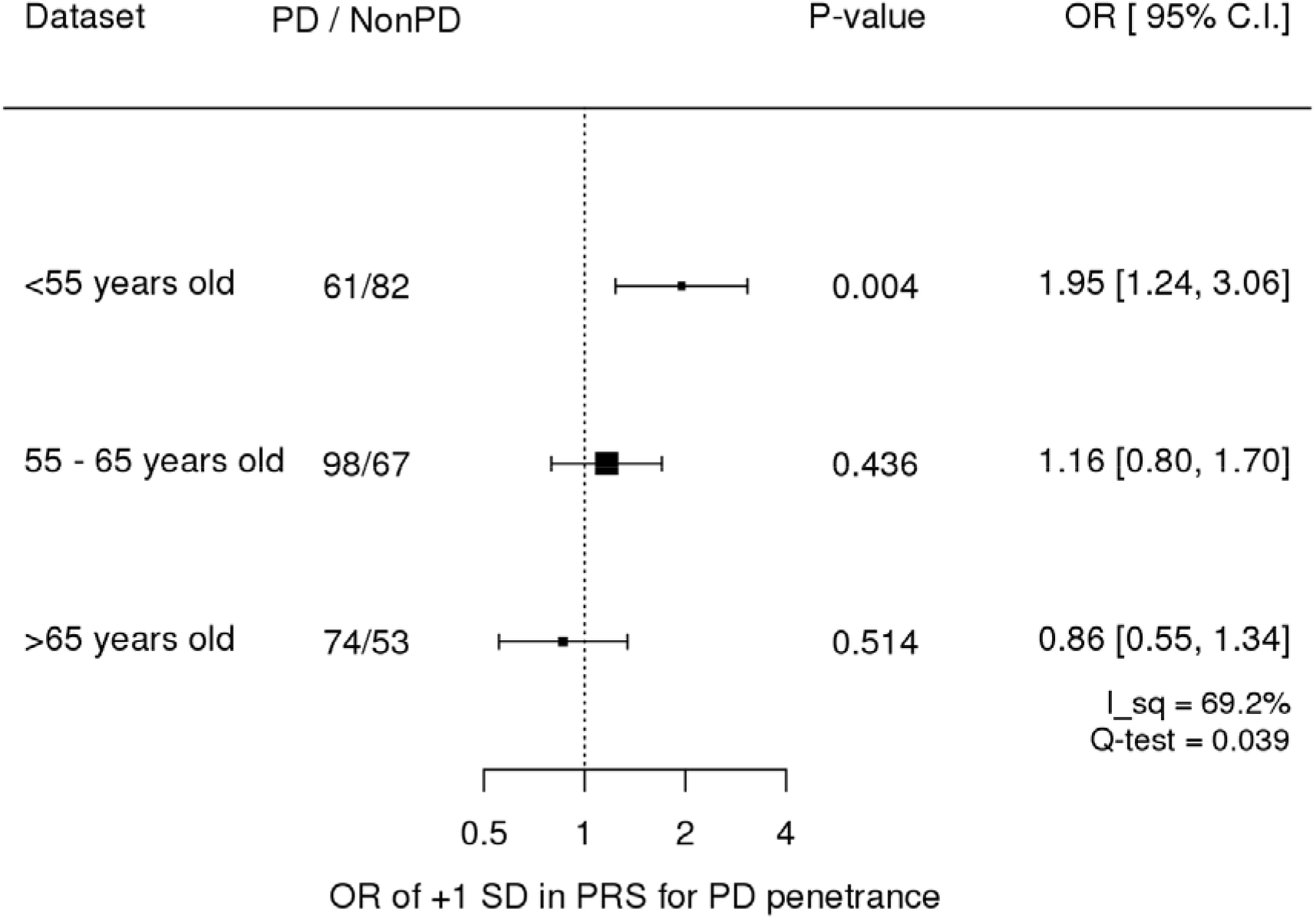
Age stratified associations of PRS on penetrance in Genetics Cohorts Dataset Legend: Age stratified analysis showed that PRS was significantly associated with the younger age. OR, Odds ratio; I_sq, I square (%); Q-test, P-value for the test of Heterogeneity. Odds ratio were adjusted for study, age, square age, sex and PC1-10

### Genetic risk score and age at onset/diagnosis in LRRK2 p.G2019S cases

Previously, we have shown that PRS is inversely correlated with age at onset in PD cases.^13^ In the current study, there was insufficient evidence that PRS was significantly associated with age at onset/diagnosis of PD in *LRRK2* p.G2019S carriers. The directionality was the same as in the general population previously studied (−0.801 [−0.959, −0.643] per +1 SD in PRS), but the association did not reach significance level (Supplemental Figure 2). As mentioned in the power calculation analyses in Methods, this is an underpowered study, and therefore additional data is needed to confirm these results.

### Exploratory analysis for the individual Parkinson’s disease-associated variants

While the power of the study is not sufficient, the motivation for this analysis was to qualitatively assess whether any of the risk variants could be strong penetrance modifiers. The Q-Q plot for the observed and expected *P* in −log10 scale showed an upper deviation from the expected line (lambda value of 1.34), indicating the contribution of these variants for the PD risk in p.G2019 LRRK2 heterozygous carriers. However, no single association stood out as a possible penetrance modifier after adjusting for multiple tests (Supplemental Figure 3). Furthermore, when the estimates for the associations of carriers and noncarriers of individual variants were compared, there were 10 variants whose tests of homogeneity were rejected at a significance level of 0.05 (Supplemental Figure 4, Supplemental Table 2). Additionally, *GBA* mutations in *LRRK2* p.G2019S carriers (Supplemental Table 3) were analyzed, but there was no clear enrichment of *GBA* mutations in cases versus controls or for younger age at onset among cases.

## Discussion

The aggregated genetic risks in the form of PRS that were constructed from the recent case-control GWAS of European population was significantly associated with the risk of PD among heterozygous *LRRK2* p.G2019S mutation carriers in our study. Furthermore, the magnitude of the association was larger in the younger population, and this interaction between age and PRS was significantly larger than in noncarriers. Conversely, the age at onset or the age at diagnosis among carriers with PD was not significantly associated with PRS, likely due to the small study size rather than a truly null association. Additionally, although age at onset and age at diagnosis is highly correlated, clinical site variation and differences in assessing this could affect the analysis. Therefore, more data are required to conclusively determine this point.

When we investigated specific variants of interest for PD, such as *GBA* coding variants and PD GWAS variants, no significant differences were identified, although relatively small numbers of *GBA* coding variants did not seem to affect *LRRK2* penetrance, which is in line with previous results.^20^ The effect size of 10 PD GWAS variants were heterogeneous between p.G2019S carriers and noncarriers at an unadjusted significance level of 0.05. We need more data to formally assess these different associations for PD between carriers and noncarriers, but genes in these loci are good candidates as potential interactors. Interestingly, rs823118 has a larger effect size and is a locus that harbors an *RAB7L1*, which is a *LRRK2* interactor.^21^ Alsp the effect size of rs1293298 (*CTSB* locus) is opposite compared to general PD, where the rs1293298 locus was recently shown to be a potential genetic modifier of *GBA*-associated PD.^15^

The primary limitation of this study is the size, despite including several large cohorts. *LRRK2* p.G2019S is rare in the general population, and therefore actively recruiting carriers would be a good strategy to increase power for studies such as ours. To investigate the penetrance of carriers, recruitment only focusing on carriers are very effective, as shown in Table 1. More initiatives and future collaboration with clinical trials only targeting *LRRK2* p.G2019S mutations are warranted. Currently, there are at least three clinical trials targeting *LRRK2*-linked PD: two use kinase inhibition, and the third focuses on reducing the protein load. As these trials move forward, important considerations are whether the disease mechanisms that cause *LRRK2*-linked PD are generalizable to typical PD, and whether it is possible to identify which *LRRK2* mutation carriers will or will not express disease. Previously, we have shown that active randomization on genetic components of PD is important for successful clinical trials because unbalanced genetic risk may lead to the heterogeneity of intervention arms and lower the power of that trial.^22^ Our results suggest that actively balancing PRS is similarly important for a disease modifying/preventing trial targeting p.G2019S carriers.

Taken together, we show that penetrance and likely age at onset of *LRRK2* is modified by PD PRS, implying a clear genetic influence.

## Acknowledgements

We would like to thank all of the subjects who donated their time and biological samples to be a part of this study. We also would like to thank all members of the International Parkinson Disease Genomics Consortium (IPDGC). See for a complete overview of members, acknowledgements and funding http://pdgenetics.org/partners. This work was supported in part by the Intramural Research Program of the National Institute on Aging, National Institutes of Health, Department of Health and Human Services; project ZO1 AG000949.

Data used in preparation of this article were obtained from the MJFF□sponsored LCC and PPMI database (www.ppmi-info.org/data). For up-to-date information on these studies, visit www.michaeljfox.org/lcc and www.ppmi-info.org. PPMI – a public-private partnership – is funded by The Michael J. Fox Foundation for Parkinson’s Research and funding partners, including AbbVie, Allergan, Avid Radiopharmaceuticals, Biogen, BioLegend, Bristol-Myers Squibb, Celgene, Denali Incorporated, GE Healthcare, Genentech, GlaxoSmithKline, Eli Lilly and Company, Lundbeck, Merck & Co., Meso Scale Discovery, Pfizer, Piramal, Prevail Therapeutics, Roche, Sanofi Genzyme, Servier Laboratories, Takeda, Teva, UCB, Verily, Voyager Therapeutics, and Golub Capital (www.ppmi-info.org/fundingpartners). LCC is coordinated and funded by The Michael J. Fox Foundation for Parkinson’s research. The investigators within the LCC contributed to the design and implementation of the LCC and/or provided data and/or collected biospecimens, but did not necessarily participate in the analysis or writing of this report. The full list of LCC investigators can be found at www.michaeljfox.org/lccinvestigators.

## Appendix

Supplemental Figure 1. PRS of case and control in IPDGC and UKBB

Supplemental Figure 2. Age at onset/diagnosis and the PRS

Supplemental Figure 3. The QQ plot for the population risk variants.

Supplemental Figure 4. Population risk variants whose effect sizes are heterogeneous between carriers and non-carriers

Supplemental Table 1. Homozygous carriers of LRRK2 p.G2019S

Supplemental Table 2. Variants whose effect on the risk are homogeneous between carriers and non-carriers

Supplemental Table 3 Contingency Table for GBA protein coding variants in LRRK2 p.G2019S carriers

IPDGC collaborator’s list

## Authors’ Role

1) Research project

1A. Conception: CB, IPDGC, AS

1B. Organization: HI, CB, IPDGC

1C. Execution;HI, CB, IPDGC

2) Statistical Analysis

2A. Design: HI, CB

2B. Execution:HI, CB

2C. Review and Critique: All

3) Manuscript:

3A. Writing of the first draft: HI, CB,

3B. Review and Critique: All

## References

1. Nalls MA, Blauwendraat C, Vallerga CL, et al. Expanding Parkinson’s disease genetics: novel risk loci, genomic context, causal insights and heritable risk. bioRxiv. Epub 2019.

2. Nichols WC, Pankratz N, Hernandez D, et al. Genetic screening for a single common LRRK2 mutation in familial Parkinson’s disease. Lancet. 2005;365:410–412.

3. Gilks WP, Abou-Sleiman PM, Gandhi S, et al. A common LRRK2 mutation in idiopathic Parkinson’s disease. Lancet. 2005;365:415–416.

4. Bras JM, Guerreiro RJ, Ribeiro MH, et al. G2019S dardarin substitution is a common cause of Parkinson’s disease in a Portuguese cohort. Mov Disord. 2005;20:1653–1655.

5. Ozelius LJ, Senthil G, Saunders-Pullman R, et al. LRRK2 G2019S as a Cause of Parkinson’s Disease in Ashkenazi Jews. N Engl J Med. 2006;354:424–425.

6. Lesage S, Dürr A, Tazir M, et al. LRRK2 G2019S as a Cause of Parkinson’s Disease in North African Arabs. N Engl J Med. 2006;354:422–423.

7. Healy DG, Falchi M, O’Sullivan SS, et al. Phenotype, genotype, and worldwide genetic penetrance of LRRK2-associated Parkinson’s disease: a case-control study. Lancet Neurol. 2008;7:583–590.

8. Lee AJ, Wang Y, Alcalay RN, et al. Penetrance estimate of LRRK2 p.G2019S mutation in individuals of non-Ashkenazi Jewish ancestry. Mov Disord. 2017;32:1432–1438.

9. Trinh J, Gustavsson EK, Vilariño-Güell C, et al. DNM3 and genetic modifiers of age of onset in LRRK2 Gly2019Ser parkinsonism: a genome-wide linkage and association study. Lancet Neurol. 2016;15:1248–1256.

10. Fernández-Santiago R, Garrido A, Infante J, et al. α-synuclein (SNCA) but not dynamin 3 (DNM3) influences age at onset of leucine-rich repeat kinase 2 (LRRK2) Parkinson’s disease in Spain. Mov Disord. 2018;33:637–641.

11. Blauwendraat C, Reed X, Kia DA, et al. Frequency of Loss of Function Variants in LRRK2 in Parkinson Disease. JAMA Neurol. 2018;75:1416.

12. Whiffin N, Armean M I, Kleinman A, et al. Human loss-of-function variants suggest that partial LRRK2 inhibition is a safe therapeutic strategy for Parkinsons disease. bioRxiv. Epub 2019.:1–34.

13. Blauwendraat C, Heilbron K, Vallerga CL, Bandres-Ciga S, Coelln R von, Pihlstrøm L. Parkinson disease age at onset GWAS: defining heritability, genetic loci and α-synuclein mechanisms. Mov Disord. 2019;in press.

14. Nalls MA, Escott-Price V, Williams NM, et al. Genetic risk and age in Parkinson’s disease: Continuum not stratum. Mov Disord. 2015;30:850–854.

15. Blauwendraat C, Reed X, Nalls M, Krohn L, Heilbron K. Genetic modifiers of risk and age at onset in GBA positive Parkinson’s disease and Lewy body dementia. preprint. Epub 2019.

16. McCarthy S, Das S, Kretzschmar W, et al. A reference panel of 64,976 haplotypes for genotype imputation. Nat Genet. 2016;48:1279–1283.

17. Das S, Forer L, Schönherr S, et al. Next-generation genotype imputation service and methods. Nat Genet. 2016;48:1284–1287.

18. Bycroft C, Freeman C, Petkova D, et al. The UK Biobank resource with deep phenotyping and genomic data. Nature. 2018;562:203–209.

19. Van Hout V C, Ioanna T, Beckman D J, et al. Whole Exome Sequencing and Characterization of Coding Variation in 49,960 Individuals in the UK Biobank. bioRxiv. Epub 2019.

20. Yahalom G, Greenbaum L, Israeli-Korn S, et al. Carriers of both GBA and LRRK2 mutations, compared to carriers of either, in Parkinson’s disease: Risk estimates and genotype-phenotype correlations. Parkinsonism Relat Disord. 2019;62:179–184.

21. Beilina A, Rudenko IN, Kaganovich A, et al. Unbiased screen for interactors of leucine-rich repeat kinase 2 supports a common pathway for sporadic and familial Parkinson disease. Proc Natl Acad Sci. 2014;111:2626–2631.

22. Leonard H, Blauwendraat C, Krohn L, et al. Genetic variability and potential effects on clinical trial outcomes: perspectives in Parkinson’s disease. bioRxiv 2018.

